# Multi-glomerular projection of single olfactory receptor neurons is conserved among amphibians

**DOI:** 10.1101/788133

**Authors:** Lukas Weiss, Lucas D. Jungblut, Andrea G. Pozzi, Barbara S. Zielinski, Lauren A. O’Connell, Thomas Hassenklöver, Ivan Manzini

## Abstract

Individual receptor neurons in the peripheral olfactory organ extend long axons into the olfactory bulb forming synapses with projection neurons in spherical neuropil regions, called glomeruli. Generally, odor map formation and odor processing in all vertebrates is based on the assumption that receptor neuron axons exclusively connect to a single glomerulus without any axonal branching. We comparatively tested this hypothesis in multiple fish and amphibian species by applying sparse cell electroporation to trace single olfactory receptor neuron axons. Sea lamprey (jawless fish) and zebrafish (bony fish) support the unbranched axon concept, with 94% of axons terminating in single glomeruli. Contrastingly, axonal projections of the axolotl (salamander) branch extensively before entering up to six distinct glomeruli. Receptor neuron axons labeled in frog species (Pipidae, Bufonidae, Hylidae and Dendrobatidae) predominantly bifurcate before entering a glomerulus and 59% and 50% connect to multiple glomeruli in larval and post-metamorphotic animals, respectively. Independent of developmental stage, lifestyle and adaptations to specific habitats, it seems to be a common feature of amphibian olfactory receptor neuron axons to frequently bifurcate and connect to multiple glomeruli. Our study challenges the unbranched axon concept as a universal vertebrate feature and it is conceivable that also later diverging vertebrates deviate from it. We propose that this unusual wiring logic evolved around the divergence of the terrestrial tetrapod lineage from its aquatic ancestors and could be the basis of an alternative way of odor processing.

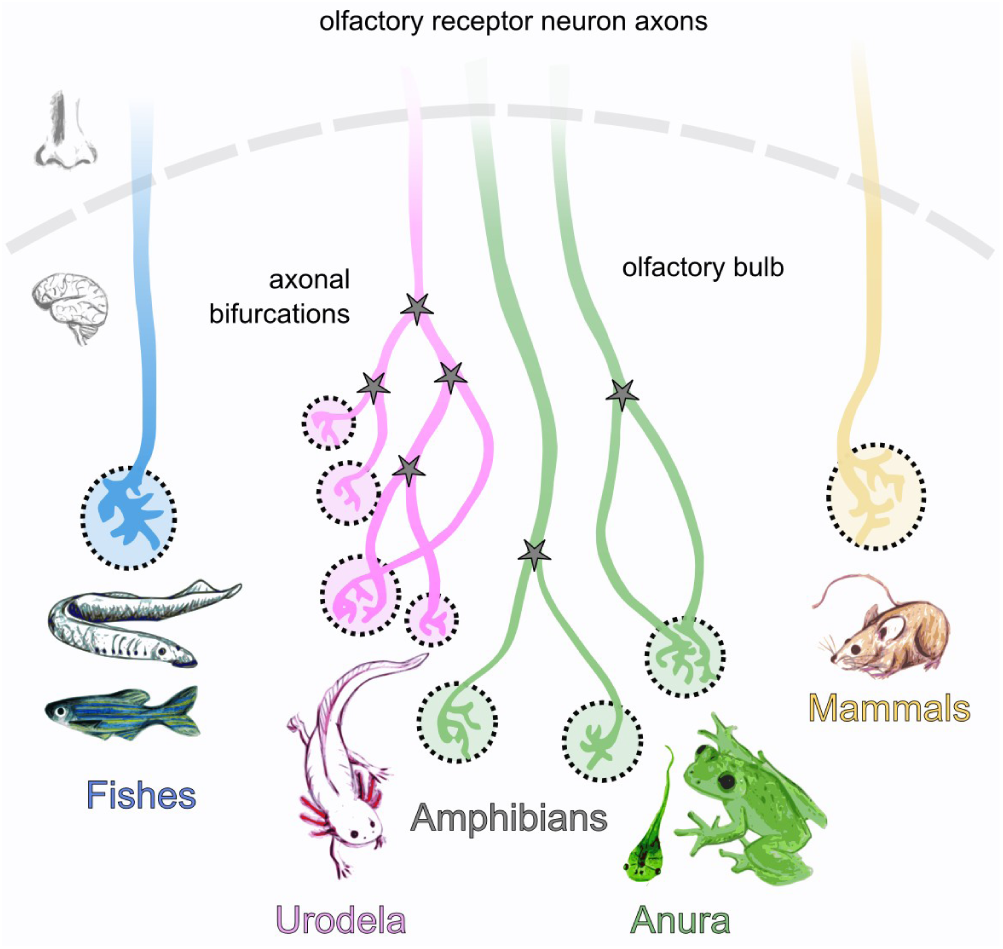

## Introduction

Vertebrates are equipped with a sophisticated olfactory system to detect relevant chemical information about their environment. Throughout vertebrate evolution, a progressive segregation into parallel olfactory pathways takes place [1–3]. While the peripheral olfactory organ of fishes consists of a single olfactory surface [4–6, but see 7], the mammalian system is segregated into several anatomically and functionally distinct subsystems [2,8]. A first bipartition of the olfactory periphery into a main olfactory epithelium (MOE) and a vomeronasal organ (VNO) coincides with the evolution of the first tetrapods, the amphibians [9]. However, primordial structures that could potentially be homologous to the VNO have been identified in earlier diverging vertebrates like lungfish [10] and lamprey [7].

Chemical detection in the various sensory epithelia is relying on the expression of olfactory receptor proteins in the dendritic cilia or microvilli of olfactory receptor neurons (ORNs) [11], with the two major receptor gene families being the OR-type olfactory receptor genes and the vomeronasal receptor genes [12,13]. In the main olfactory system of rodents, each ORN expresses a single OR-type olfactory receptor [14,15] and sends a single, unbranched axon to the olfactory bulb (OB) via the olfactory nerve (ON). In the main OB, an axon terminally branches in the confines of a single dense neuropil structure, a glomerulus. All axons of ORNs equipped with the same olfactory receptor type coalesce onto one or very few glomeruli [16–18]. Each glomerulus in the main OB is thus believed to relay the information of a single OR-type olfactory receptor to the postsynaptic projection neurons (PNs), distinguished as mitral and tufted cells in rodents [1]. This constitutes the idea of the chemotopic organization of the rodent main OB [19]. While mammalian PNs extend their single primary dendrite into one sole glomerulus, PNs in fish, amphibians and reptiles often bear several primary dendrites connecting to multiple glomeruli [for review see 20].

In contrast to the wiring logic employed by the main olfactory system, all vomeronasal receptor neurons (VRN) in the VNO expressing the same type of vomeronasal receptor converge onto ∼15-30 glomeruli in the mammalian accessory olfactory bulb (AOB) [21,22]. PNs in the AOB of many animals have long been known to extend multiple dendrites into spatially distinct glomeruli [23]. However, it is still widely unclear whether the synaptic input that a single PN in the AOB receives from multiple glomeruli contains the information conveyed by different or by the same type of vomeronasal receptor [24,25].

The basic olfactory wiring principles have long been assumed to be uniform among vertebrates. The first vertebrate species that has been found to violate the rule of an unbranched ORN axon innervating a single glomerulus in the main OB was the African clawed frog *Xenopus laevis* [26]. The majority of examined axons were shown to be connecting to more than one glomerulus in larval animals [26] and this alternative pattern was retained after metamorphosis [27]. It remains elusive whether this multi-glomerular wiring is a specific adaptation of the secondarily aquatic *Xenopus* or if it is a more conserved evolutionary feature also present in other vertebrate lineages.

Here, we report that bifurcating ORN axons and multi-glomerular innervation are not a particular adaptation of *Xenopus laevis*, but a conserved feature throughout the order Anura (frogs and toads). ORN axon tracings in four ecologically diverse frog species in pre- and post-metamorphotic animals showed that this alternative olfactory wiring scheme is independent of developmental stage and of habitat. We could also show that multi-glomerular innervation of single ORN axons are the predominant pattern in the axolotl salamander, which suggests that this feature might be present in all amphibians. Contrastingly, both the main olfactory system of the sea lamprey (jawless fish) as well as the olfactory system of zebrafish (teleost fish) follow the unbranched ORN axon paradigm with a single ORN axon only arborizing within the confines of a single glomerulus. We propose that the unusual wiring logic found in amphibians evolved around the divergence of the terrestrial tetrapod lineage from its aquatic ancestors and forms the basis of an alternative way of odor processing.

## Results

### ORN axons in fish have less branching points than amphibian axons

The morphology of an ORN axon in vertebrates is generally described as an unbranched projection terminating in fine arborizations within a single glomerulus of the OB [28]. It was already reported that this principle does not apply to the wiring scheme in the secondarily aquatic African clawed frog [26,27]. We investigated whether this alternative projection pattern could be more common in the olfactory system of other aquatic vertebrates. We traced single ORNs from the olfactory epithelium of the fully aquatic post-larval sea lamprey (*P. marinus*, jawless fish), the zebrafish (*D. rerio*, bony fish), the axolotl (*A. mexicanum*, urodela) and the larval clawed frog (*X. tropicalis*, anura; phylogenetic overview in Figure 1B) to their axon terminals in the glomeruli of the OB. No experiments were conducted in the accessory olfactory system.

**Figure 1:**
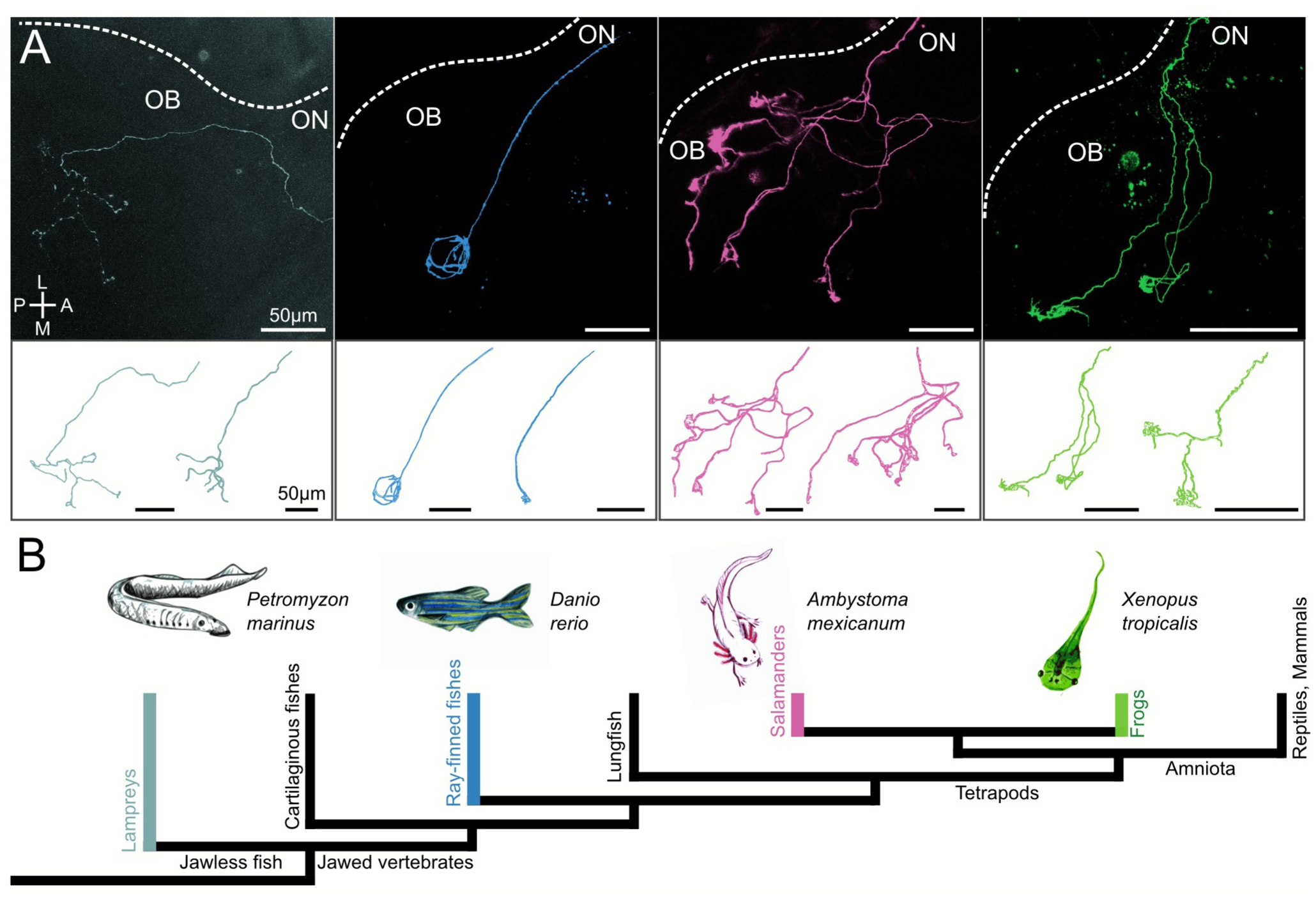
ORN axons in the OB of different aquatic vertebrates. **A)** Single ORN axons of the juvenile sea lamprey (*P. marinus;* grey), zebrafish (*D. rerio;* blue), axolotl (*A. mexicanum;* magenta) and larval clawed frog (*X. tropicalis;* green) show different levels of branching complexity. Sea lamprey and zebrafish axons are unbranched until they reach their terminals and only have a limited number of subbranches. Axolotl and frog axons branch in proximity to the ON and exhibit more subbranches. Axon tracings are shown from the transition between ON and OB until their terminals (upper panel). Dotted white line indicates the outline of the OB. The lower panel shows representative 3D reconstructions of two axons for each species. The first reconstruction of each species depicts the ORN axon shown in the upper panel. **B)** The four examined species cover a broad evolutionary period from the divergence of the jawed vertebrates from their jawless ancestors to the emergence of the first tetrapods. All four species lead a fully aquatic lifestyle. OB olfactory bulb, ON, olfactory nerve, ORN olfactory receptor neuron, P posterior, A anterior, L lateral, M medial.

Axon tracings of the four species differed substantially in their general branching structure (Figure 1A). ORN axons in the OB of the sea lamprey and the zebrafish showed similarity with the pattern reported for rodents [28]. A long, unbranched axon projects towards the glomerular layer of the OB where it terminally arborizes. While both species follow this common feature, the zebrafish axons have even shorter and fewer terminal arborizations than the lamprey axons. In contrast, the ORN axons of the two amphibian species bifurcate shortly after entering the OB, projecting several sub-branches into the glomerular layer, where each sub-branch arborizes again. Quantifying the bifurcations of each single axon along the distance from its entry point in the OB to the axon terminals, we found significant differences between fishes and amphibians (Figure 2A and B). Sea lamprey axons have on average 6.5 ± 4.5 terminal arborizations (n = 6). This pattern is not significantly different from the zebrafish axons that show even fewer arborizations (3 ± 1.4, n = 10). Zebrafish axons were found to have the least complex structure, with the highest amount of arborizations for a single axon being five. The axolotl displays a significantly different pattern from both the lamprey and zebrafish axons (p = 0.0058 and 0.00001 respectively, Figure 2B), the ten examined axons bifurcate 29.7 ± 7.7 times on average, with one axon even branching 42 times. Axonal tracings obtained from the tadpoles of the clawed frog (23 ± 8.1, n = 10) also showed a significantly higher degree of branching when compared to the zebrafish (p = 0.0013). They did not show significantly different branching points than lamprey (p = 0.08) or axolotl axons.

**Figure 2:**
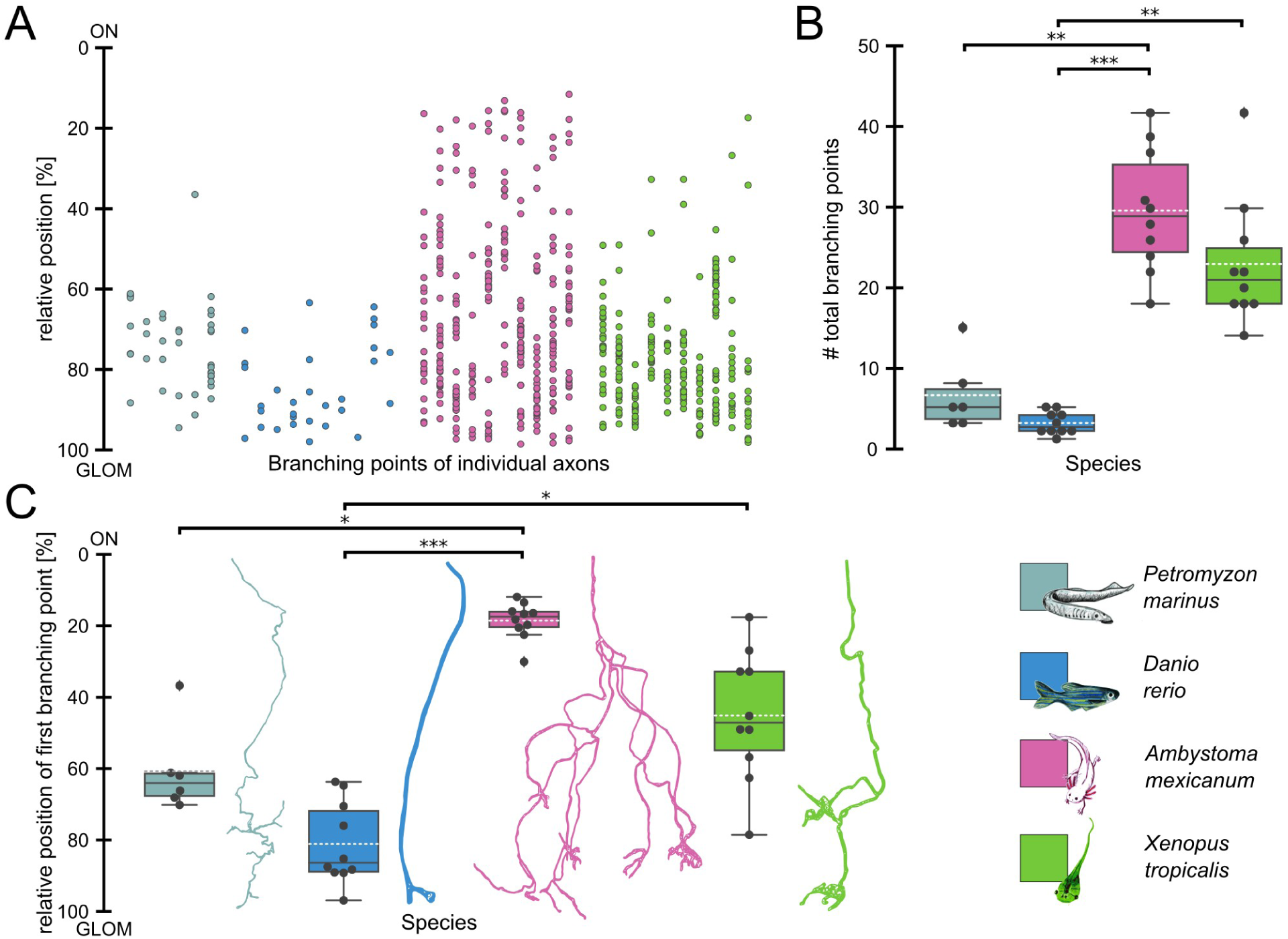
Number of axonal branching points and relative position of the first axonal bifurcation differ in fishes and amphibians. **A)** Branching points of individual ORN axons and their relative position between the transition from ON to OB (0%) and their most distal axon terminal in the glomerular layer of the OB (100%) are shown. All dots on a vertical line depict the positions of all branching points of a single reconstructed axon. ORN axons of sea lamprey (n = 6; grey), zebrafish (n = 10; blue), axolotl (n = 10, magenta) and of the clawed frog (n = 10; green) are shown. **B)** Quantitative comparison of the total amount of ORN axonal branching points in each species. Each dot represents a single ORN axon. The black line indicates the median, the white dotted line the mean amount of branching points for axons of each species. Axolotl ORN axons have significantly more branching points than lamprey (p = 0.0058) and zebrafish axons (p = 0.00001). The ORN axons of the western clawed frog are significantly more branched than the zebrafish axons (p = 0.0013). **C)** Species comparison of the first axonal bifurcation of ORN axons. The relative position of the first bifurcation between the transition from ON to OB (0%) and their most distal axon terminal (100%) are shown. The relative position of the first branching point is closer to the axon terminals in the glomerular layer in the OB of both fish species. The axolotl axons branch in immediate proximity of the ON, significantly different from the fish axons (lamprey, p =0.014 and zebrafish p = 0.000001). Frog axons also branch closer to the ON, significantly different from the zebrafish (p = 0.014). A representative axonal reconstruction is shown for each species. Statistical significance was tested using Kruskal-Wallis rank sum test followed by Dunn’s multiple comparison post-hoc test with Holm-Bonferroni correction. OB olfactory bulb, ON olfactory nerve, ORN olfactory receptor neuron, GLOM glomerular layer.

In addition to the amount, also the spatial distribution of branching points seems to be differently organized in fishes and amphibians (Figure 2C). The point of origin for distance measurements was set as the transition between ON and OB. While most fish ORN axonal projections only arborize close to their terminals in the glomerular layer, amphibian axons start to bifurcate much closer to the ON and in the nerve layer (Figure 2C). The first lamprey axon bifurcation happens around 359 ± 139 µm after entering the OB, which measures 61 ± 12% of the distance from the origin to the furthest terminal point. The average unbranched axon segment of the zebrafish tracings was 184 ± 39 µm (81 ± 12%), axolotl axons first branch at 85 ± 33 µm (18 ± 5%) and the larval clawed frog axons at 94 ± 40 µm (45 ± 18%). There is a significant difference between the relative position of the first bifurcation in axolotl axons and axons of both fish species (lamprey: p = 0.014, zebrafish: p = 0.000001). The position of the first bifurcations in *Xenopus* axons are not statistically different from the ones in lamprey or axolotl axons, but different from zebrafish (p = 0.014, Figure 2C). There are substantial differences in the general ORN axon architecture between the four examined species. Our results clearly indicate that ORN axon bifurcations before reaching the glomerular layer cannot be attributed to an aquatic habitat, since this feature is absent both in the sea lamprey and the zebrafish. In contrast, our tracings suggest that this alternative branching pattern of bifurcations rostral to the glomerular layer could be linked more specifically to the amphibian lineage, since it was found to be present in both a salamander and a frog species.

### Multi-glomerular ORN axons are present only in amphibians

While it was reported for the rodent main olfactory system that single ORN axons project into a single glomerulus [17,18], 86% of ORNs in the main OB of larval *X. laevis* were connected to more than one glomerulus [26]. So far, comparative data of other vertebrate species on this matter are missing. To classify uni- and multi-glomerular ORN axons in the various animal species, we used an algorithm identifying glomerular clusters based on the spatial density of branching and endpoints of the reconstructed axonal structures.

We found that out of all fish axons (lamprey: n = 6, zebrafish: n = 10), only one zebrafish axon was classified as multi-glomerular, while the remaining axons projected into a single target structure and are thus considered uni-glomerular (Figure 3A and B). Only three fish axons (one lamprey axon and two zebrafish axons) had a single extra-glomerular branching point (average extra-glomerular branching points lamprey: 0.2 ± 0.4, zebrafish: 0.2 ± 0.4), while all other axons only started to arborize within the target glomerular cluster (Figure 3C). The average distance from the transition between ON and OB to the glomeruli measured 401 ± 111 µm in sea lamprey and 194 ± 41 µm in zebrafish axons. In the axolotl, all except for one axon (n = 10) followed a multi-glomerular output pattern (Figure 3A and B), with four axons even innervating six distinct glomerular structures. The average axonal distance from the entry point in the OB to the glomeruli was 336 ± 109 µm. Axolotl axons bifurcate 5.1 ± 1.9 times on average before reaching their target glomeruli (Figure 3C), which is significantly different from both lamprey and zebrafish (p = 0.0002 and 0.00001 respectively).

**Figure 3:**
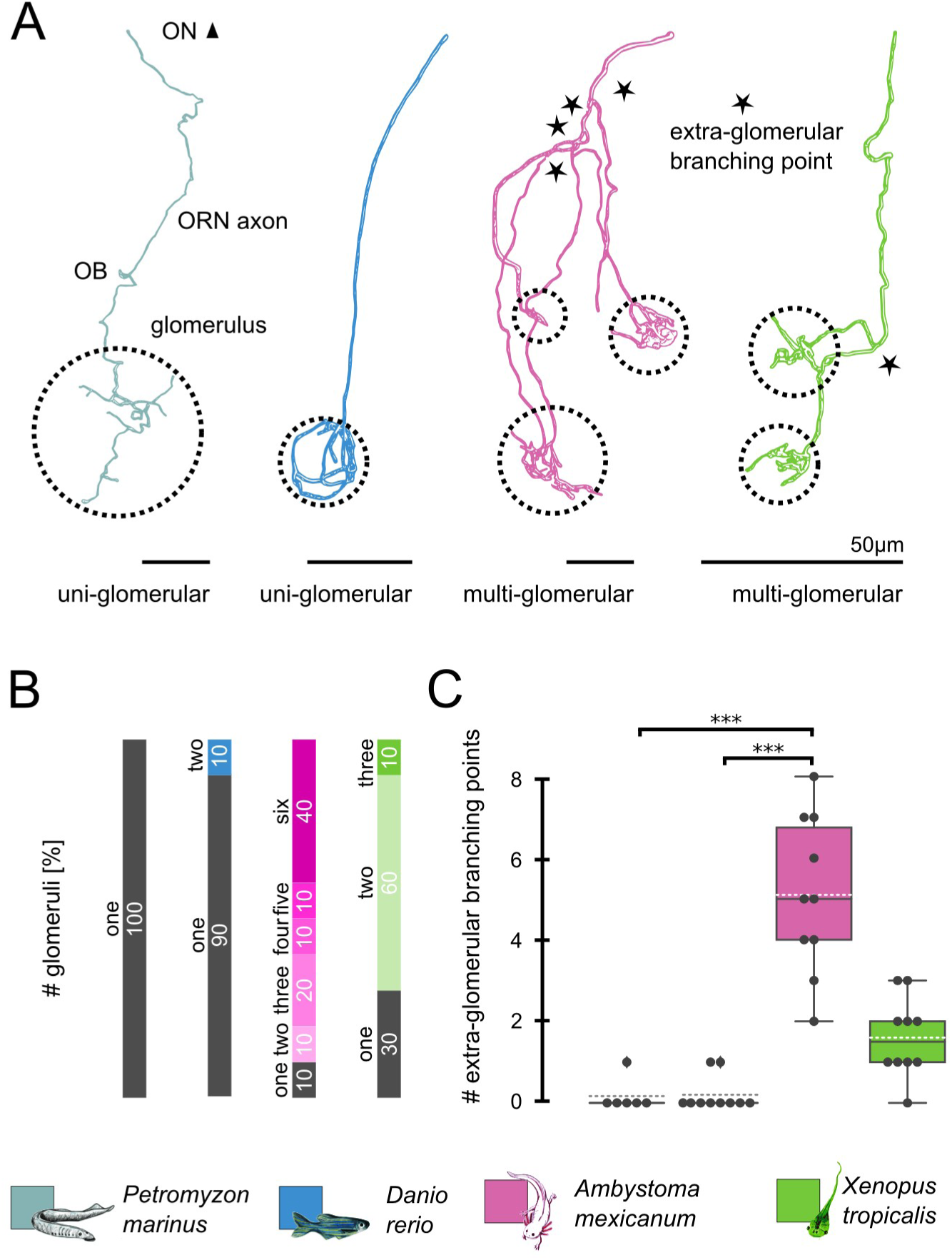
Lamprey and zebrafish ORN axons connect to a single glomerulus, amphibian ORN axons are multi-glomerular. **A)** The representative ORN axonal reconstructions of the lamprey (grey) and the zebrafish (blue) both connect to a single glomerulus (dotted circle) and have no bifurcations prior to entering the glomerulus. The representative axolotl ORN axon (magenta) branches four times (black stars) before connecting to three glomeruli, the frog axon (green) has a single extra-glomerular branching point and innervates two separate glomeruli. **B)** Population analysis of all examined axons, each stacked bar represents the percentual share of uni- and multi-glomerular axons (two to six glomeruli) of one of the four species. All lamprey ORN axons (n = 6) and 90% (n = 10) of zebrafish axons were classified as uni-glomerular by the DBSCAN algorithm. On the contrary, most amphibian axons were classified as multi-glomerular (axolotl 90%, clawed frog 70%), with a varying number of innervated glomeruli. **C)** Quantitative comparison of the amount of extra-glomerular branching points of all axons of each species. Each dot represents a single ORN axon. The black line indicates the median, the white dotted line the mean amount of extra-glomerular branching points. Axolotl axons have a significantly higher amount of extra-glomerular branching points than lamprey (p = 0.0002) and zebrafish (p = 0.00001). Statistical significance was tested using Kruskal-Wallis rank sum test followed by Dunn’s multiple comparison post-hoc test with Holm-Bonferroni correction. OB olfactory bulb, ON, olfactory nerve, ORN olfactory receptor neuron. The clawed frog *Xenopus tropicalis* belongs to the Pipidae family and is classified as a Mesobatrachian, an evolutionarily more basal frog species. The other three species belong to the evolutionarily more ‘modern’ frogs, the Neobatrachians, and to the families of Bufonidae, Hylidae and Dendrobatidae. The diverse ecology of both the adult frogs (above) and their larval offspring (below) are summarized next to the drawings. The tree is pruned from the anuran tree in [48], which originally included 3309 species.

The anuran olfactory system (represented by the clawed frog), showed a more heterogeneous wiring pattern. Three out of ten tracings were classified as uni-glomerular, while seven axons were multi-glomerular. In comparison with the axolotl, *Xenopus* axons projected into maximally three glomeruli, while the majority (n = 6) displayed a bi-glomerular wiring pattern (Figure 3A and B). The mean axonal distance from the nerve to the glomeruli was 162 ± 43 µm, the structures branched 1.6 ± 1.0 times before ending in glomerular clusters (Figure 3C). The branching density inside a single glomerular cluster was not significantly different between lamprey (6.3 ± 4.7), zebrafish (2.6 ± 1.1) and axolotl (8.5 ± 7.5). *Xenopus* glomeruli were more densely packed with sub-branches (12.8 ± 4.8) compared to zebrafish (p = 0.00009), which displayed the lowest bifurcation density, often with only one branching point within a glomerulus. Our results support the hypothesis that multi-glomerular innervation of single ORN axons is not a wiring feature exclusive to *X. laevis* but seems to be common among other aquatic amphibian species of both salamanders and frogs. On the other hand, uni-glomerular wiring is the prevalent - almost exclusive - pattern in fishes, which strongly resembles the rodent wiring logic.

### The alternative wiring pattern in anurans is independent of developmental stage and ecology

The alternative wiring pattern of multi-glomerular ORN axons has so far only been described in amphibians that live a water-bound lifestyle [27]. While all amphibian larvae are dependent on an aquatic habitat, most adult frogs leave the water after metamorphosis [29]. We conducted ORN tracing experiments in anurans with more or less water-independent adult lifestyles to test whether the alternative olfactory wiring is an olfactory adaptation of aquatic frogs and tadpoles or habitat independent. To account for the diverse ecology of the anurans, we examined the following species in addition to the aquatic *X. tropicalis*. Larval *Rhinella arenarum* are vegetarian grazers and adults are terrestrial. Both *Scinax granulatus* and *Ranitomeya imitator* have carnivorous/omnivorous tadpoles, their adults are arboreal and terrestrial, respectively. The major difference between these two species is that *Ranitomeya* provides extensive parental care and the number of tadpoles per parent pair is much lower than in *Scinax, Rhinella* and *Xenopus* (Figure 4) [29].

**Figure 4:**
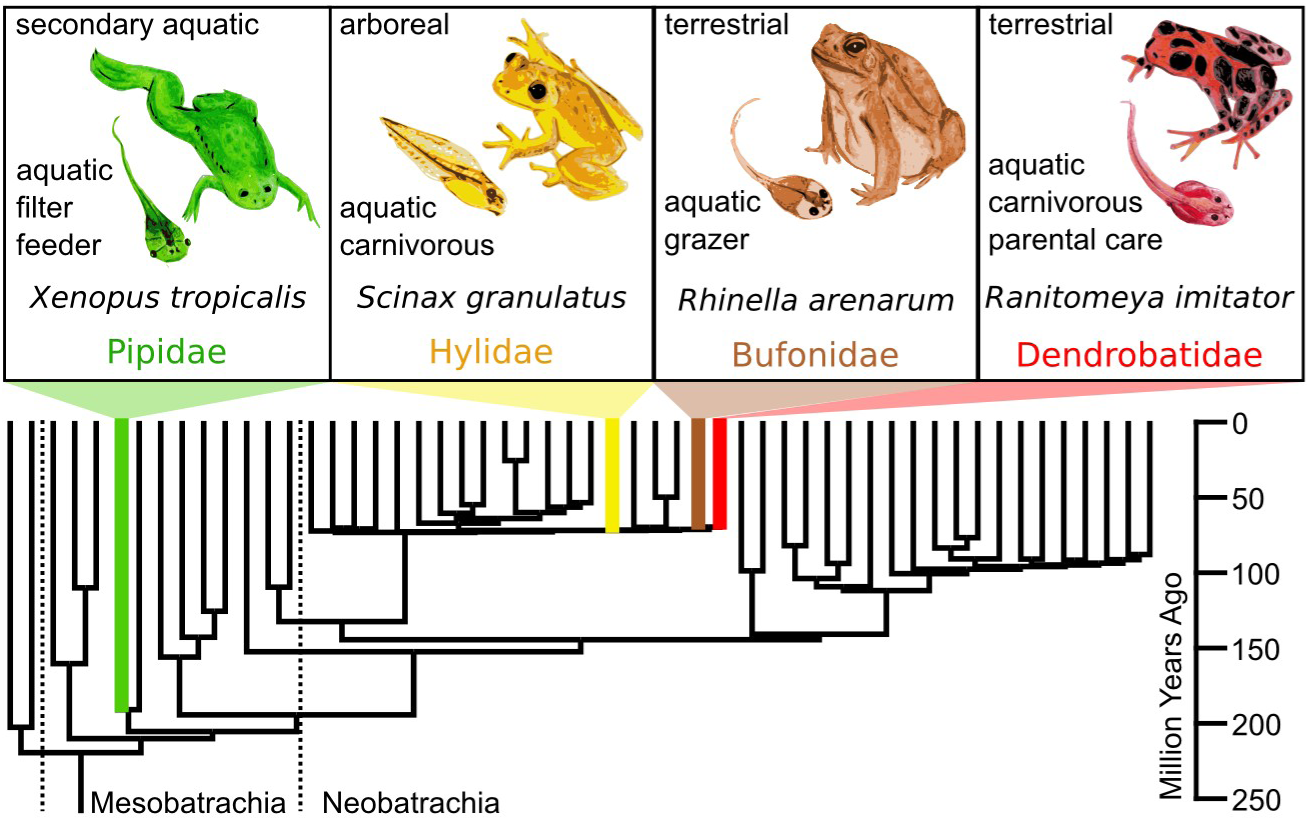
Overview about ecology and phylogeny of the four anuran species examined in this study.

The tracings of ORN axons in *R. arenarum, S. granulatus* and *R. imitator* tadpoles showed similar wiring and branching properties to those found in *X. tropicalis* tadpoles. In Figure 5A two representative axonal reconstructions per species are shown. White stars indicate extra-glomerular bifurcations and white dotted circles highlight glomerular clusters. The number of branching points prior to entering the glomeruli was similar in all four species (*X. tropicalis*: 1.6 ± 1.0, n = 10, *R. arenarum*: 2.4 ± 1.8, n = 8, *S. granulatus*: 1.4 ± 1.2, n = 8, *R. imitator*: 1.5 ± 1.6, n = 6), as was the distance of the first branching point relative to the entire length of the axon from the transition ON-OB to the glomeruli (*X. tropicalis*: 45 ± 18%, *R. arenarum*: 43 ± 26%, *S. granulatus*: 41 ± 28%, *R. imitator*: 45 ± 19).

**Figure 5:**
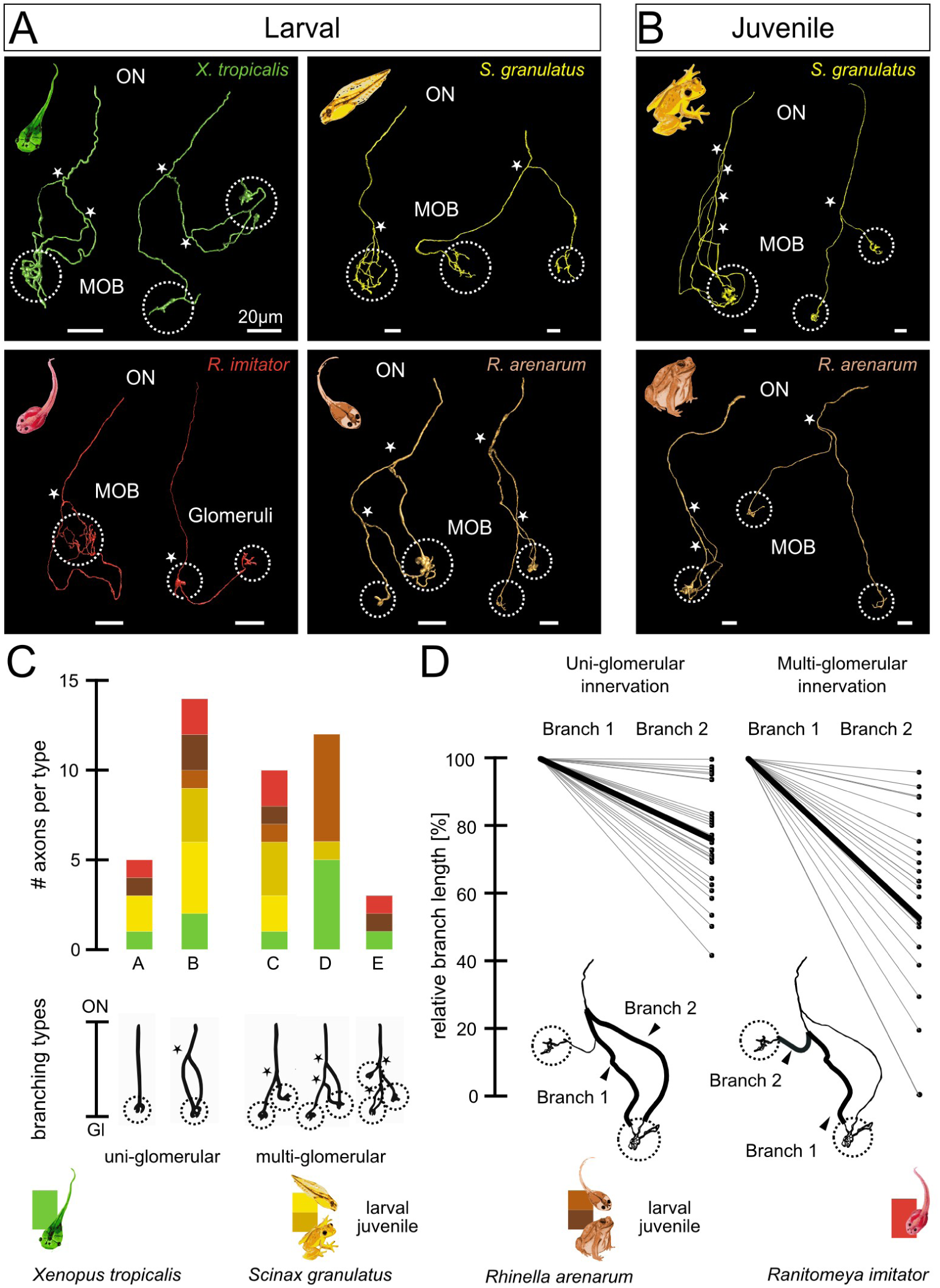
Comparison of the ORN axon morphologies in different larval and juvenile frog species with different lifestyles. **A)** Two representative ORN axon reconstructions are shown for tadpoles of each of the anuran species *X. tropicalis, R. arenarum, R. imitator* and *S. granulatus*. Each axon was reconstructed starting from the transition between ON and main OB to the most distal axon terminal. Multiple axonal branches innervate one or two glomeruli (white dotted circles), and generally have one or two extra-glomerular branching sites (white stars). **B)** Two representative ORN axon reconstructions are shown for post-metamorphotic terrestrial juveniles of *R. arenarum* and arboreal *S. granulatus.* Axonal structures are comparable to their larval counterparts shown in A, one or two glomeruli are innervated by two or more axonal subbranches. **C)** ORN axons were classified into five branching types (A and B uni-glomerular, C - E multi-glomerular). Types are schematically shown below, black stars indicate extra-glomerular branching points. The number of axons belonging to each branching type is shown in the stacked bar plot above. We included all ORN axons of tadpoles (*X. tropicalis*, green, n = 10; *S. granulatus*, light yellow, n = 8; *R. arenarum*, light brown, n = 8; *R. imitator*, red, n = 6) and juveniles (*S. granulatus*, dark yellow, n = 7; *R. arenarum*, dark brown, n = 5). The counts of axons per type for each species and developmental stage are marked according to the color legend below. **D)** Relative length of the longer and the shorter branch of individual ORN axons innervating the same glomerulus (left plot) or two different glomeruli (right plot). Each pair of dots refers to a single axon, where branch 1 is the longer branch and branch 2 is the shorter one. The thick black line visualizes the mean length relation between the two branches. In uni-glomerular axons, the shorter branch measures 76% of the length of the longer branch (n = 27, ORN axons of all species and developmental stages included). In multi-glomerular axons, the shorter branch measures on average 53% the length of the longer branch innervating a different glomerulus (n = 25, ORN axons of all species and developmental stages included). MOB main olfactory bulb, ON, olfactory nerve, ORN olfactory receptor neuron.

To exclude that the alternative multi-glomerular wiring pattern is linked to the larval stages and/or their aquatic lifestyle, we conducted sparse cell electroporation in juveniles of the terrestrial *R. arenarum* and the arboreal *S. granulatus*. Figure 5B shows reconstructions of single ORN axons of juveniles of the two species. Their morphology does not significantly differ from the morphology of conspecific tadpoles nor from the other larval axons examined. Before reaching the glomeruli, axons of juvenile *R. arenarum* and *S. granulatus* bifurcate 2.2 ± 1.5 (n = 5) and 1.6 ± 1.1 (n = 7), respectively. The first bifurcations occur at 48 ± 29% (*R. arenarum*) and 28 ± 16% (*S. granulatus*) of the total distance between the ON-OB transition and the axon terminals. The only parameter that slightly differs from axons of their larval conspecifics (other than an approx. 1.5-fold increase in total axonal length from tadpoles to juveniles) is the amount of branching points inside the glomerular clusters (*R. arenarum*: larval 5.7 ± 1.4, juvenile 3.8 ± 0.9, *S. granulatus*: larval 11.7 ± 3.1, juvenile 5.5 ± 4).

All four anuran species and ontogenetic stages were heterogeneous with regard to the number of glomerular clusters that are innervated by a single axon and the proportion between uni- and multi-glomerular axons. We categorized the axonal structures based on recurring branching patterns and classified them into five categories (type A to E; Figure 5C and D). Among the uni-glomerular axons, we distinguished two main types. Type A is characterized by a single, unbranched axon terminating in a single glomerular cluster. This is the prevailing type reported in rodents and also found in the fish species we examined here. Type B also terminates in a single glomerulus, but has at least two separate branches projecting into the same glomerular structure. Of all the axons reconstructed from larval and juvenile anurans (n = 44), only 11% belonged to Type A. Type B was more frequent, amounting to 32% of the axons. 57% of the axons were classified as multi-glomerular, with 50% of all axons innervating two glomeruli and only three axons (7%) innervating more than two glomerular end-structures (Type E). In 23% of all axons two glomerular clusters were innervated by a single branch each (Type C), in 27 % of the tracings, at least one of the two glomeruli was innervated by more than one sub-branch (Type D, Figure 5C). Among the axons traced in the various species, all species displayed at least four out of the five different types.

We additionally measured the differences in the length of branches entering the same or different glomeruli of a single axonal structures for all the anuran species. We found 27 cases (20 in larvae, 7 in juveniles) where a single glomerulus was innervated by at least two separate axonal branches (Figure 5D, left plot). By subtracting the shortest innervating branch from the longest, we measured a branch length difference of 46 ± 71 µm in tadpoles and 56 ± 27 µm in juveniles. On average, the shortest branch had 76 ± 16% the length of the longer one. This value was consistent between tadpoles (75 ± 16%) and juveniles (80 ± 12%). In all multi-glomerular axons (n = 25, larvae 19, juveniles 6), we measured the branch length difference between the branches innervating the nearest and the furthest glomerulus (Figure 5D, right plot). In larval anurans, the difference was 58 ± 46 µm, in juveniles 154 ± 214 µm. The distance to the glomerulus closer to the nerve layer was 53 ± 30 % of the distance to the glomerulus most distant from the nerve layer (tadpoles: 48 ± 31%; juveniles: 67 ± 25%).

The collected data indicates that ORN axonal projections in larval and juvenile amphibians are much more heterogeneous than what has been reported in rodents and what we found in fishes. In amphibians, multi-glomerular innervation is retained throughout their developmental stages and does not seem to be linked to a specific lifestyle or habitat.

### Emergence of the bifurcation of ORN axon coincides with the first tetrapods

In an attempt to put our findings into an evolutionary context, we compared the wiring properties of the fishes and urodela to the anuran juveniles, to give an overview about the presumably mature system of the animals. Since the axolotl is a neotenic salamander, the main olfactory system of the one to two month old larva is assumed to be a mostly developed system that does not undergo drastic changes until sexual maturity is reached. The majority of the fish axons (75%) are uni-glomerular and unbranched prior to entering the glomeruli (Figure 6, blue bars), which is in accordance with the prevailing vertebrate wiring principle. The juvenile anurans are the most heterogeneous group. Only the minority of axons follow the unbranched axon principle (8%). Most axons branch at least once before terminating in one or more glomeruli. 42% of the axons innervate one glomerulus with more than two separate branches, 50% innervate more than two glomeruli (Figure 6, yellow bars). Of the salamander axons, 80% innervate more than three glomerular structures (Figure 6, magenta bars). None of the axolotl axons followed the unbranched one axon-one glomerulus principle.

**Figure 6:**
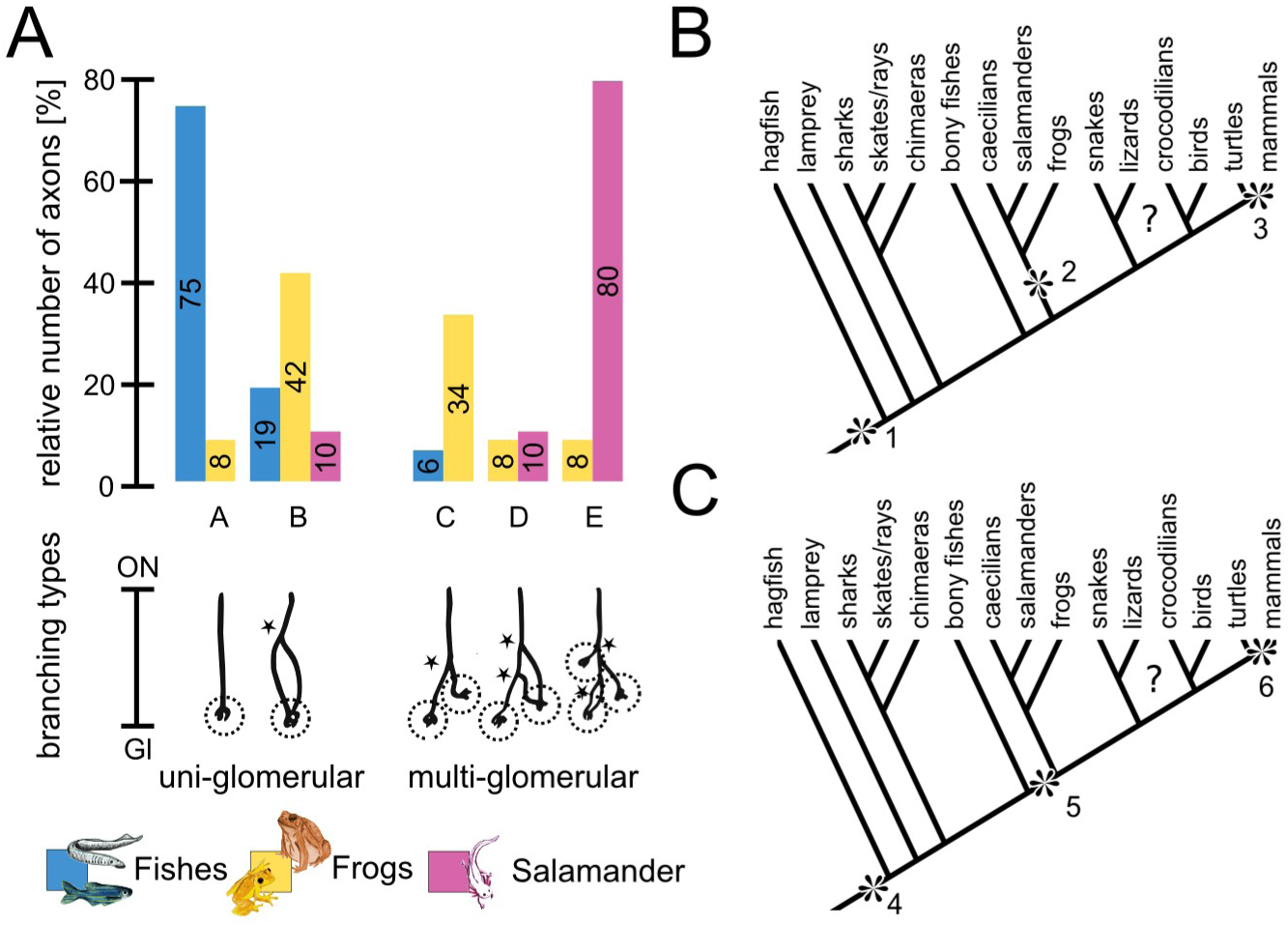
Multi-glomerular innervation pattern is a conserved feature of amphibians and could have emerged with the evolution of the first tetrapods. **A)** ORN axons of fish species (lamprey and zebrafish, blue bars, n = 16), juvenile frogs (*R. arenarum and S. granulatus*, yellow bars, n = 12) and the axolotl salamander (magenta bars, n = 10) were classified into five branching types, (A and B uni-glomerular, C-E multi-glomerular). Types are schematically shown, black stars indicate extra-glomerular branching points, dotted circles indicate glomeruli. The relative number of axons belonging to each branching type for each group is shown in the bar plots. The majority of fish axons follow the unbranched axon pattern (type A) and the majority of salamander ORN axons innervate three or more glomeruli (type E). Frog axons are more heterogeneous with the most represented types being bi-glomerular (type C) and uni-glomerular with at least one extra-glomerular branching point (type B). **B)** One possible evolutionary scenario for the emergence of the multi-glomerular ORN projections could be that the unbranched, uni-glomerular ORN axon is a basic vertebrate trait (*1) and has secondarily evolved into a multi-glomerular alternative wiring logic in the amphibian lineage (*2) and independently also in the accessory system of mammals (*3). **C)** In a second scenario, the basic unbranched ORN wiring logic (*4) changed to the alternative bifurcating axon logic in vertebrates at the transition from an aquatic to a terrestrial habitat (*5). In mammals, this alternative logic then segregated to the accessory system, while the main olfactory system again followed the unbranched axon wiring logic (*6). The phylogenetic tree was modified from [9].

Our results strongly imply an alternative principle of odor processing on the level of the OB and that the emergence of this principle coincides with the divergence of the first terrestrial tetrapods from their aquatic ancestors. Even though data from the smallest order of modern amphibians (Caecilians) is still elusive, axonal bifurcations and multi-glomerular innervation seem to be conserved in all amphibians regardless of their developmental stage and habitat.

## Discussion

### Multi-glomerular innervation is absent in fishes and present in all developmental stages of amphibians

The consensual hypothesis regarding olfactory wiring in the vertebrate main olfactory system is established on the idea that all ORNs expressing one allele of the olfactory receptor gene repertoire transmit information into only one or two glomeruli in the main OB via an unbranched axon. This leads to the formation of a precise odotopic map, where each glomerulus is part of a unique olfactory unit that encodes the information detected by a specific receptor type [16–18]. In this study we show that ORN axons of post-larval sea lamprey and zebrafish clearly follow the unbranched axon principle postulated for all vertebrates [16–18]. Fish axons solely arborize within their target-glomerulus and only one axon out of 16 has been found to innervate two distinct glomeruli in the zebrafish. In contrast, 57% of ORN axons labeled in anurans show connections to multiple glomeruli (90% in the salamander species examined).

It is still under debate how the precise wiring between ORN axons and PN dendrites within the glomeruli is established and whether there are extensive pruning mechanisms taking place during development. Experiments in newborn rabbits [30] and mice [31,32] have shown that exuberant ORN axon growth in the main OB during early development is subsequently pruned. Contrastingly, other studies conducted in neonatal rats [28] and zebrafish embryos [33] show that ORN axons arborize to their final morphology without erroneous targeting. Tenne-Brown and colleagues demonstrate the occurrence (15% of all axons) of extra-glomerular bifurcations and multi-glomerular connections of single ORN axons in neonatal mice, but only until two weeks after birth [32]. Of the above mentioned studies, only Klenoff and colleagues report the occurrence of axons arborizing in two multiple glomeruli in rats after the initial pruning phase is over, yet only as a very rare exceptions (<0.1% of axons) [28].

In contrast, we found that multi-glomerular wiring is preserved even after metamorphosis is finished in all examined amphibians. 50% of the axons analyzed in juveniles of *R. arenarum* and *S. granulatus* innervated more than one glomerulus and 92% of the axons bifurcated before entering the glomerular structures. A study conducted in the main OB and the AOB of larval and post-metamorphotic *Xenopus laevis* [27] has yielded very similar results, showing that neither pre-glomerular bifurcations nor multi-glomerular innervation can be solely attributed to the larval stages. However, in accordance with the results shown by Marcucci and colleagues [31], we noticed a reduction of the total number of branching points from larval to post-metamorphotic animals. Even though the number of extra-glomerular bifurcations (larval 1.7 ± 1.4, n = 32; juvenile 1.8 ± 1.3, n = 12) remained constant, the arborizations inside a single glomerulus decreased in the juveniles (larval 10.1 ± 5.1; juvenile 4.8 ± 3.1). While the reduction of branches inside a glomerulus can be explained by pruning mechanisms during development, multi-glomerular innervation in the amphibian main OB is not strictly linked to an immature larval stage.

### The alternative wiring logic in amphibians has parallels to the rodent AOB

In the rodent accessory olfactory system, the axons of VRNs expressing the same vomeronasal receptor type converge onto ∼15-30 glomeruli, constituting a more vague spatial code than in the main OB [13,21,22]. In contrast to the main system, the postsynaptic partners of the ORN axons, the PNs, extend several dendrites into multiple glomeruli. It is still unclear whether they integrate between input of VRNs expressing different or the same vomeronasal receptors [21,22,24,25]. Several attempts have been made to unravel the AOB wiring logic from VRN axons to PN dendrites in rodents. A study investigating genetically labeled receptor neurons expressing a single type of V1R or V2R in mice gives evidence for a homotypic connectivity model. In this model, a single PN extends its dendrites into multiple glomeruli innervated exclusively by the same vomeronasal receptor type [24]. Another study supported a selective heterotypic connectivity model in which a single PN receives information from glomeruli that get sensory input not from a single vomeronasal receptor type, but from closely related receptors within a receptor subfamily [25].

The discrepancies between the wiring principles in the rodent main OB and AOB suggest that there might be different aspects of odor information extracted by the respective subsystems. It has additionally been reported that approx. 10% of single VRN axons in the nerve layer of the AOB split into several sub-branches, reaching out to multiple glomeruli [34]. The wiring pattern of amphibians described in our study therefore resembles the rodent accessory system rather than the main system. While the rodent system shows a clear separation into the OR-type receptor expressing MOE and the V1R and V2R expressing VNO, this segregation is incomplete in amphibians. Among the OR-type receptors and other receptor gene families, the MOE of *Xenopus* was shown to express V1Rs as well as early diverging V2Rs, while the *Xenopus* VNO is expressing more recently diverging V2R genes [1,35,36].

In line with this heterogeneous receptor gene expression in the amphibian MOE, we found quite a heterogeneity in ORN axon branching patterns. Different branching types could be connected to different receptors. The mix of ORs and VRs in the MOE of amphibians could explain the heterogeneity of branching patterns, while in the more segregated rodent system, axonal bifurcations and a more vague spatial glomerular code are only found in the accessory system. It has already been shown that olfactory receptors are involved in axon guidance and the formation of the glomerular map [19,37]. Still, the expression of vomeronasal receptors in the MOE of amphibians is unlikely to be the only cause for axonal bifurcations and the alternative wiring pattern. V1Rs and V2Rs are already expressed in the sensory epithelium of fishes, but we could show that ORN axon bifurcations are absent in both lampreys and zebrafish. It is more plausible that the mechanisms by which the receptors influence axonal guidance could have changed over evolutionary time and that the new wiring principle has only emerged after the divergence of the first tetrapods from the aquatic ancestors.

### Multi-glomerular ORN innervation is mirrored by PN morphology in vertebrates and invertebrates

Just like the PNs in the rodent AOB, PNs in the main OB of amphibians and reptiles extend multiple dendrites into multiple glomeruli, where they terminate in dendritic tufts [20]. In the case of amphibians, the multi-glomerular PN morphology seems to be mirroring the multi-glomerular ORN axons described in this study. In the sea lamprey, the morphology of the single ORN axons we found show a net like branching structure within single glomeruli. This morphology is also mirrored in the uni-glomerular arborizations of the lamprey PNs [38]. In many teleost fish species, it is known that PNs extend many primary dendrites into multiple glomeruli. However, this was only shown to be the minority in zebrafish [39]. In accordance with these results, we also show that single zebrafish ORN axons mostly (90%) terminate in a single glomerulus. It is intriguing to speculate, whether teleost fishes equipped with multi-glomerular PNs also display multi-glomerular ORN wiring.

A similar mirror image in the connectivity pattern between ORN axon and postsynaptic PNs occurs in the evolution of the antennal lobe in orthopteran insects [40]. It was shown that in more basal orthopterans (*e.g*. the great green bush cricket), single receptor neuron axons are innervating a single glomerulus and a single PN extends its dendrite into one sole glomerulus, resembling the mammalian main system. Contrastingly, in later diverging orthopterans (locusts and grasshoppers) both single receptor neurons as well as PNs connect to multiple glomeruli – a pattern similar to amphibians [40–42]. The multi-glomerular pattern in locusts is linked to the formation of a high number of microglomeruli (∼2500 glomeruli), while the one-to-one pattern in basal orthopterans is linked to fewer number of bigger glomeruli (∼40) [41]. A similar evolution towards microglomeruli could have taken place among vertebrates: the sea lamprey has very large but few glomeruli (41-65) [43], the zebrafish has approx. 140 quite differently sized glomeruli [44] and *Xenopus laevis* has a larger number of smaller glomeruli, ∼350 in the main OB [45] and ∼340 in the AOB [46]. The concept of the branched and multi-glomerular ORN axon has thus developed at least twice independently, however its putative functional implications remain elusive.

### Different evolutionary scenarios for the emergence of the alternative wiring logic

Branched receptor neuron axons with multi-glomerular innervation have been shown in about 10% of VRNs in the mouse AOB [34] and as a predominant type in the main OB [26] and the AOB of *Xenopus laevis* tadpoles, as well as adults [27]. In this study we show that this wiring logic is also present in the axolotl salamander and in anuran tadpoles of four ecologically distinct families (Pipidae, Bufonidae, Hylidae and Dendrobatidae) as well as terrestrial post-metamorphotic frogs. From our results, we can conclude that the multi-glomerular ORN wiring pattern is independent of tadpole or adult ecology and seems to be a feature derived from the common ancestor between frogs and salamanders, since it is also present in axolotl. In both the jawless and bony fish species we examined, we could not find any clear signs of the presence of this alternative wiring principle.

From an evolutionary perspective, our results suggest several plausible scenarios, two of which will be discussed here. First, the axonal bifurcations of ORNs could have developed independently in amphibians (*2, Fig 6B), and in the VNO of mammals (*3, Fig 5B), with the ancestral vertebrate trait being the one-to-one wiring logic (*1, Fig 5B). The presence of multi-glomerular wiring structures in the orthoptera supports the idea that this alternative wiring could have a functional advantage in odor processing and has thus evolved independently multiple times [40]. In the second scenario, the multi-glomerular pattern could have evolved around the divergence of the terrestrial tetrapod lineage from its aquatic ancestors (*5, Fig 5B), coinciding with the first occurrence of the VNO [9] and a huge expansion and reshaping of the OR-type gene family [1,47]. Given that the odorant receptor is directly influencing axon targeting and the formation of glomeruli [16,37], it could potentially also have an impact on axonal branching. Newly emerging receptors or axon branching principles after the tetrapod divergence could be responsible for this alternative wiring and be more widespread in the MOE and VNO of earlier diverging tetrapods (i.e. amphibians), and more focally expressed in the VNO of mammals (*6, Fig 5B).

Taken together, our results show that the prevailing idea of an unbranched ORN axon arborizing only in a single glomerulus cannot be generalized for vertebrates any longer. While jawless and bony fishes display the one-to-one wiring logic that is shown for the rodent main olfactory system, our results indicate that bifurcating ORN axons and multi-glomerular wiring are a general feature of the amphibian olfactory system. This alternative wiring scheme is neither linked to larval stages, nor to a specific habitat or lifestyle of amphibians. It cannot be excluded that this feature is even more common amongst vertebrates and that it constitutes the basis of an alternative way for odor processing.

## Acknowledgements

We thank all the present and past members of the Manzini laboratory for fruitful discussion and input, especially Thomas Offner and Sara Joy Hawkins. We thank Anja Schnecko for dedicated animal care and Gianfranco Grande and Eva Fischer for support to set up experiments. L.W. was granted a travel fellowship (JEBTF-180809) by the company of Biologists Limited and the Journal of Experimental Biology to visit the O’Connell lab. This work was supported by DFG Grant 4113/4-1 and in part, by Award Number S10RR02557401 from the National Center for Research Resources (NCRR). Its contents are solely the responsibility of the authors and do not necessarily represent the official views of the NCRR or the National Institutes of Health.

## Author Contributions

Conceptualization, L.W., T.H. and I.M.; Investigation, Formal Analysis, Visualization and Writing – Original Draft, L.W.; Writing – Review & Editing, L.W., L.D.J., A.G.P., B.S.Z., L.A.O., T.H. and I.M.; Funding Acquisition and Resources, L.D.J., A.G.P., B.S.Z., L.A.O., T.H. and I.M.; Supervision, T.H. and I.M.

## Declaration of Interest

The authors declare no competing interests.

## Experimental Procedures

### 1. Animals

#### Fish species

All sea lampreys (*Petromyzon marinus*) used in this study were post-larval transformer stages (metamorphic stage seven, both sexes) from the Connecticut River, Turner Falls, MA. Animals were captured and supplied by United States Geological Survey Conte Anadromous Fish Research Laboratory. They were kept in 420 l tanks at 6°C ± 1°C under static renewal conditions until used. Zebrafish (*Danio rerio*, both sexes) were kept in oxygenated water tanks at room temperature.

#### Amphibian species

Albino larvae of axolotl (*Ambystoma mexicanum*) were obtained from the Ambystoma Genetic Stock Center at the University of Kentucky, USA. They were kept in oxygenated water tanks (20°C) and fed with red mosquito larvae. Animals used for this study were both sexes, 5-6 weeks of age. Wild type *Xenopus tropicalis* larvae were bred and reared at the Institute of Animal Physiology, University of Giessen. They were kept in water tanks at a water temperature of 25°C and fed with algae. Animals used for this study were stages 49-52 after Niewkoop and Faber [49].

*Ranitomeya imitator* tadpoles of stages 28-31 after Gosner [50] were bred in the laboratory colony at the Biology Department of Stanford University, Palo Alto, CA, USA. Individual tadpoles were kept separately after hatching at a water temperature of 25°C.

*Rhinella arenarum* larvae (Gosner stages 29-31) and juvenile animals were obtained by *in vitro* fertilization from a colony at the Faculdad de Ciencias Exactas y Naturales of the University of Buenos Aires. *Scinax granulatus* larvae (Gosner stages 31-35) and juveniles were collected from the wild (semi-temporary ponds formed in the surroundings of the Campus of the University of Buenos Aires). All larval animals were kept in tanks of dechlorinated water at a temperature of 22°C and fed ad libitum with chard leaves. Juvenile animals were kept in glass terraria and fed with flies.

All experiments performed followed the guidelines of Laboratory animal research of the Institutional Care and Use Committee of the University of Windsor (AUPP 14-05), University of Buenos Aires (CD: 316/12, Protocol #22), the University of Göttingen (33.9-42502-04-12/0779), University of Gießen, (GI 15/7, 932_GP) and Stanford University (APLAC-33016).

### 2. Sparse cell electroporation

Animals were anesthetized using 0.02% MS-222 (ethyl 3-aminobenzoate methanesulfonate; Sigma-Aldrich) in tap water until completely unresponsive and placed on a wet tissue paper under a stereomicroscope. ORNs were stained using micropipettes pulled from borosilicate glass capillaries (Warner instruments, resistance 10-15 MΩ) filled with fluorophore-coupled dextrans (Alexa dextran 488 and 594, 10 kDa, Life Technologies) diluted at a concentration of 3mM in saline Ringer (Amphibian Ringer (mM): 98 NaCl, 2 KCl, 1 CaCl_2_, 2 MgCl_2_, 5 glucose, 5 Na-pyruvate, 10 Hepes, pH 7.8; Lamprey Ringer (mM): 130 NaCl, 2.1 KCl, 2.6 CaCl_2_, 1.8 MgCl_2_, 4 Hepes, 4 dextrose, 1 NaHCO_3_, pH 7.4, Zebrafish Ringer (mM): 131 NaCl, 2 KCl, 20 NaHCO_3_, 1.25 KH_2_PO_4_, 2.5 CaCl_2_, 2 MgSO_4_, 10 dextrose, 5 Na-pyruvate, 10 Hepes, pH 7.2). The pipettes were mounted on the electrode bearing headstage of an Axoporator 800A (Axon instruments, Molecular Devices) and inserted into the main nasal cavity of the animals. A 500 ms train of square voltage pulses (50 V, single pulses 300 µs at 200–300 Hz) was triggered to stain neurons [27,51]. This protocol was repeated at different positions inside the main nasal epithelia. The animals were left to recover.

### 3. Olfactory bulb whole mount preparation

Animals were anesthetized again (as described above) three days after electroporation and killed by severing the spinal cord at the level of the brainstem. The whole olfactory bulbs were dissected out of the tissue. Samples were immediately imaged in Ringer’s solution or fixed in 4% PFA in PBS for one hour and imaged later.

### 4. Image acquisition and processing

Multi-channel image stacks (z-resolution of 1 µm) were acquired using multi-photon microscopy at an excitation laser-wavelength of 780 nm (upright Nikon A1R-MP and upright Leica SP5 multiphoton microscopes). Brightness and contrast of the stacks were adjusted using ImageJ [52]. Since there was no dye introduced into the tissue with a blue-wavelength emission, we used the blue-wavelength detector (400 – 492 nm) to image tissue auto-fluorescence. Pigmentation derived autofluorescence was mathematically subtracted from the other emission channels using the image-calculator function implemented in ImageJ. Image data are presented as maximum intensity projections along the z-axis of the virtual image stacks.

### 5. Axonal reconstructions

Individual neuronal morphology was reconstructed semi-automatically by defining branching and endpoints of the axonal structure from the acquired image stacks in Vaa3D [53]. Only ORN axons that could be traced from the beginning of the nerve layer in the olfactory bulb until their terminals in the glomerular layer were reconstructed. The reconstructed neuron-trees were sorted by the sort_swc algorithm implemented in Vaa3D to define the root of the axon as first node of the structure.

### 6. Structural analysis and identification of glomeruli

Axonal reconstructions were analyzed and quantified using custom written Python scripts in Jupyter notebook. Spatial distribution of branching points and length of branches and sub-branches were assessed based on the reconstructions. All data presented in regard to branch length is measured in distances along the axonal structure (in µm) or displayed as length-ratio in %.

The number of glomeruli innervated by an axonal structure was determined using the DBSCAN algorithm (Density-Based Spatial Clustering of Applications with Noise) implemented in the sklearn machine learning package written for Python [54]. The algorithm clusters together points that are in close spatial proximity to a lot of neighboring points (glomerular cluster) while it marks points far away from its closest neighbors as low-density noise. All branching- and endpoints of a neuron-tree structure from the root to the terminals of the axons were used as input points. Based on the algorithm, an axon terminal was considered a glomerular cluster if at least three points were in close spatial proximity. Blunt axonal endings without terminal bifurcations were marked as noise outliers. To account for the different glomerular size between the various animals used for this study, the minimal distance for points to be considered a cluster was chosen separately for each species and ontogenic stage (in µm: *P. marinus* 90, *D. rerio* 20, *A. mexicanum* 20, *X. tropicalis* 11, *R. arenarum* pre-metamorphotic 10, post-metamorphotic 15, *S. granulatus* pre-metamorphotic 14, post-metamorphotic 16, *R. imitator* 10). These values were chosen based on previously reported glomerular size (*P. marinus:* [55]; *D. rerio:* [44] or our own glomerular tracing experiments (all amphibian species).

### 7. Statistical Analysis

Averaged data are presented as mean ± standard deviation. Statistical significance was determined by Kruskal-Wallis rank sum test followed by Dunn’s multiple comparison post-hoc test, unless otherwise stated. To control familywise error rate for multiple comparisons, a Holm-Bonferroni correction was applied.

